# Gravity highlights a dual role of the insula in internal models

**DOI:** 10.1101/659870

**Authors:** Célia Rousseau, Marie Barbiero, Thierry Pozzo, Charalambos Papaxanthis, Olivier White

**Affiliations:** INSERM UMR1093-CAPS, Université Bourgogne Franche-Comté, UFR des Sciences du Sport, F-21000, Dijon; IIT@UniFe Center for Translational Neurophysiology of Speech and Communication, Istituto Italiano di Tecnologia, Via Fossato di Mortara, 17-19, Ferrara, Italy

**Keywords:** motor imagery, graviception, internal simulation, arm movements

## Abstract

Movements rely on a mixture of predictive and reactive mechanisms. With experience, the brain builds internal representations of actions in different contexts. Many factors are taken into account in this process among which the immutable presence of gravity. Any displacement of a massive body in the gravitational field generates forces and torques that must be predicted and compensated by appropriate motor commands. Studies have shown that the insular cortex is a key brain area for graviception. However, none attempted to address whether the same internal representation of gravity is shared between reactive and predictive mechanisms. Here, participants either mentally simulated (only predictive) or performed (predictive and reactive) vertical movements of the hand. We found that the posterior part of the insular cortex was engaged when feedback was processed. The anterior insula, however, was activated only in mental simulation of the action. A psychophysical experiment shows participants’ ability to integrate the effects of gravity. Our results demonstrate a dual internal representation of gravity within the insula and discuss how they can conceptually be linked.

## Introduction

Mental simulation is a precious tool for the human mind to recall previous or to anticipate future events. Notably, through mental actions, we can simulate the consequences of our movements on the environment without physically interacting with it. This mental process is particularly beneficial when physical movements are not possible; for example, for a patient in a bed rest. It is captivating that mental and actual actions engage similar neural networks, such as the parietal and prefrontal cortices, the supplementary motor area, the premotor and primary motor cortices, the basal ganglia, the cerebellum, and even the spinal cord (Grosprêtre et al., 2016; Guillot and Collet, 2005; Hardwick et al., 2018; Hétu et al., 2013; Jeannerod, 2001). At the computational level, evidence support the hypothesis that mental simulation of movement is generated by internal forward models, which are neural networks that mimic the causal flow of the physical process by predicting the future sensorimotor state (e.g., position, velocity) given the efferent copy of the motor command and the current state (Kilteni et al., 2018; Miall and Wolpert, 1996; Wolpert and Flanagan, 2001). This computational perspective assumes that actual and mental movements trigger similar motor representations (simulation Theory, Jeannerod, 2001) and similar predicted sensory consequences (emulation theory, Grush, 2004). Interestingly, motor imagery circumvents any influence of sensory feedback: while mental movement is only a prior of motor planning, actual movement embeds motor planning corrected by sensory feedback. In both cases, sensorimotor information about the initial state is available. Contrasting these techniques offer an approach to probe the influence of feedback on an action.

Up to now, most studies have investigated mental actions by analysing kinematic variables. One interesting question, therefore, is whether dynamical parameters are also reflected into the mental movement simulation. It is well established that the brain stores internal representations of physical laws to efficiently control movements in a challenging environment (Angelaki et al., 2004; Barbiero et al., 2017; Gaveau et al., 2016; Mcintyre et al., 2001; Wolpert and Ghahramani, 2000). One of the most omnipresent and constant environmental features is gravitational force. The neural representation of gravity optimizes movement execution (Berret et al., 2008; Crevecoeur et al., 2009; Gaveau et al., 2016; White et al., 2008), interaction with falling objects (Lacquaniti et al., 2013; Zago et al., 2008; Zago and Lacquaniti, 2005), and body perception (Angelaki et al., 2004, 1999; Laurens et al., 2013; Merfeld et al., 1999), by solving the ambiguity between gravity and inertial acceleration (Einstein’s equivalence principle). Previous studies showed that the interaction with visual objects relies on an internal model of gravity, stored in the vestibular cortex. This internal representation of gravity is, however, only activated by visual motion that appears to be coherent with natural gravity (Indovina et al., 2005; Lacquaniti et al., 2014). Interestingly, the same authors found that mental imagery of objects’ visual motion does not have access to the internal model of Earth gravity, but resorts to a simulation of visual motion compatible with a 0g environment (Gravano et al., 2017). Therefore, it seems that the effects of gravitational acceleration on falling objects are considered only when one is physically interacting with the external environment. A remaining question, however, is whether an internal model of gravity is triggered when we mentally simulate our own body movements that rely on non-trivial interactions between voluntary limb acceleration and gravitational acceleration.

At the neural level, recent studies have clearly shown that, apart motor-related cortical and sub-cortical networks, the insula is a crucial brain area involved in the production of arm movements in the gravitational field (Rousseau et al., 2016). This is, however, a nuanced story. Indeed, the posterior insula is more activated only when the task involves real movements or uses visual gravity cues to plan interceptive actions (Indovina et al., 2005). In contrast, when the task is imagined (that is, only simulated without execution and feedback) or is governed by more abstract rules, the neural activity seems more prominent in the anterior insula (Mutschler et al., 2009). Here, we use actual and mental arm movements to dissociate the roles of the anterior and posterior insulae with respect to gravity. Participants performed or imagined vertical right-hand movements during fMRI sessions. We posit that the posterior insula will be activated when a movement occurs, but not when forming a mental image of the same task. This would demonstrate a dual internal representation of gravity within the insula, but with distinct roles.

## Methods

### Brain imaging experiment

#### Participants

Twenty-six healthy adults (11 females and 15 males, mean age: 28.3±7.4 years) volunteered for the experiment. All were right handed, as assessed by the Edinburgh Handedness Inventory (Oldfield, 1971), and none of them had history of neurological disorders or any indication against a fMRI examination. The Movement Imagery Questionnaire-Revised (MIQ-RS, Loison et al., 2013) was used to evaluate participants’ motor imagery ability prior to fMRI. The average score (40±7.8; maximum score= 56) indicated good imagery ability. The entire experiment complied with the Declaration of Helsinki and informed consents were obtained from all participants. The protocol was approved by the clinical Ethics Committee of the University Hospital of Dijon (registered number 2009-A00646-51).

#### Data acquisition

All experimental sessions were conducted between 2pm and 6pm. Data were acquired using a 3T Magnetom Trio system (Siemens AG, Munich, Germany), equipped with a standard head coil configuration. We used standard single shot echo planar (EPI) T2*-weighted sequence in order to measure blood oxygenation level-dependent (BOLD) contrast. The whole brain was covered in 40 adjacent interlaced axial slices (3mm thickness, TR=3050ms, TE=45ms, flip angle=90deg), each of which was acquired within a 64×64 Matrix (FOV was 20×20 cm), resulting in a voxel size of 3.125×3.125 mm.

#### Experimental procedure

We adopted a block design paradigm that alternated periods of rest (10 volumes) and periods of either motor execution (10 volumes) or kinaesthetic motor imagery (10 volumes). In the MRI scanner, participants’ upper right limb was slightly elevated by small cushions. Their hand was in supine position (palm up) and their fingers released (Figure 1A). In the *rest* condition, participants were instructed to remain quiet, motionless, and to keep their eyes open without thinking of anything in particular. In the *executed* condition, participants performed hand flexion-extension in the sagittal plane at free pace. In the *imagined* condition, they internally simulated the same vertical hand movements (Figure 1B) without actually performing them and by adopting the same hand configuration as during the *rest* condition. Specifically, participants had to feel themselves performing hand flexions and extensions in a first-person perspective (as in Demougeot and Papaxanthis, 2011).After the *imagined* session, we debriefed participants about the quality of imagined movements. They reported the vividness of the imagined hand movements on a 7-point scale (1 corresponding to “Very difficult to feel/reproduce the motor task” and 7 “Very easy to feel/reproduce the motor task”). The average score (5±1.1) indicated that all participants were actively engaged in the motor imagery process without experiencing any difficulty. Participants executed or imagined the motor task between a “GO” and a “STOP” signal delivered by the experimenter through headphones. We repeated the *rest* and *executed* conditions or the *rest* and *imagined* conditions four times during one recording session. Therefore, 80 volumes in each experimental condition were recorded per participant (4×10 volumes in the *rest* condition and 4×x10 volumes of either the *executed* or the *imagined* condition). Half of the participants started with the *executed* condition and half with the *imagined* condition.

**Figure 1.**
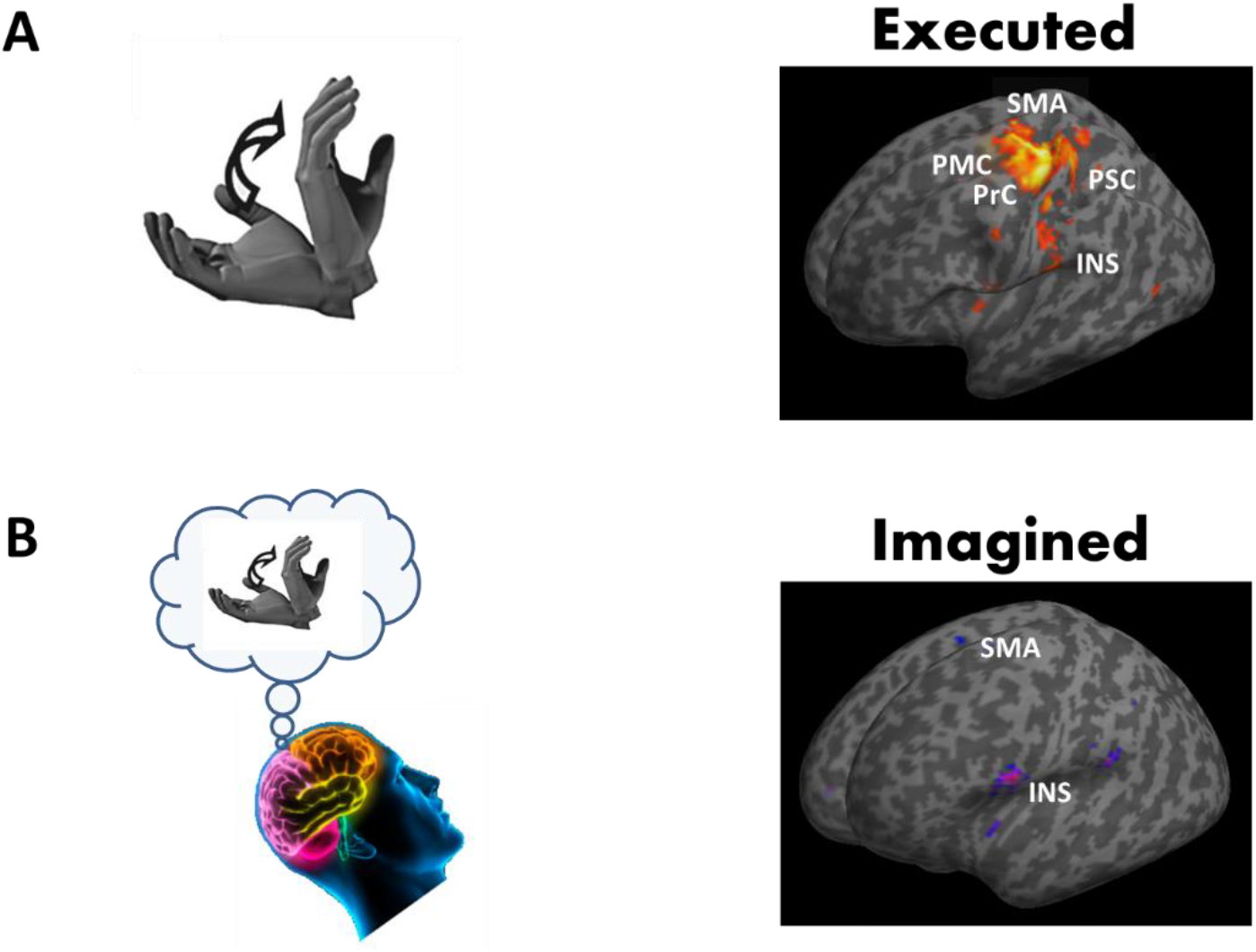
Brain areas activated during motor execution (A) and during motor imagery of hand movements (B) compared to rest. Brain responses are projected onto the inflated T1 template of MNI. Most significant brain responses are highlighted in the Primary Motor Cortex (PMC) and the Primary Somatosensory Cortices (PSC), in the Premotor Cortex (PrC) and the Supplementary Motor Area (SMA), and the Insular cortex (INS).

#### Data pre-processing and analysis

Data analysis was performed using SPM 12 (www.fil.ion.ucl.ac.uk/spm). For functional data pre-processing, each volume of both sessions was spatially realigned with the first volume of the first session using a 6-parameter fixed body transformation. Then, the T1-weighted anatomical volume was co-registered to mean images created by the realignment procedure and was normalized to the Montreal Neurological Institute (MNI) space resampled to 2 mm isotropic voxel size. The anatomical normalization parameters were subsequently used for the normalization of functional volumes. Finally, the normalized functional images were spatially smoothed with 8×8×8mm^3^ full-width at half-maximum isotropic Gaussian kernel. Time series at each voxel for each participant were high-pass filtered at 128s to remove low-frequency drifts in signal and pre-whitened by means of an autoregressive model AR(1). Data were subsequently analysed by applying a General Linear Model (GLM) separately for each participant. Blocks of *executed, imagined* and *rest* conditions were modelled using a box-car function convolved with the hemodynamic response function. Movement parameters derived from realignment corrections were also used as regressors of no interest.

At the individual level, we first assessed the whole network of brain areas involved in the processing of *executed* and *imagined* hand movements by contrasting the active phases with the *rest* blocks. Then, we performed a group analysis and applied one sample *t-tests* for the basic contrasts *executed*>*rest* and *imagined*>*rest*. A particular emphasis was given to insular cortex activation. We specifically tested insular cortex activation by contrasting *executed* and *imagined* conditions (*executed*-*rest*>*imagined*-*rest* and *imagined*-*rest*> *executed*-*rest*). We used an anatomical mask of the bilateral insula when performing group analysis and applied *one sample t-tests* for each of the two contrasts. For the whole brain network analysis, as well as for the insular cortex analysis, clusters of activated voxels were identified based on the intensity of the individual response for all contrasts (p<0.05, corrected for multiple comparisons with Bonferroni correction, t>6.3). An extended threshold of 10 voxels was determined empirically and then used for all contrasts. Results of brain activations were characterized in terms of their peak height and spatial extent and were presented in normalized stereotactic space (MNI). Brain responses were identified by means of the anatomic automatic labelling (Tzourio-Mazoyer et al., 2002).

### Psychophysical experiment

#### Participants

Fifteen right-handed healthy adults (6 females and 9 males, mean age: 25.3±7.8 years), who did not participate in the fMRI experiment, volunteered for the psychophysical experiment. All participants had normal or corrected to normal vision and none of them had history of neurological disorders, neuromuscular or chronic disease. The Movement Imagery Questionnaire-Revised (MIQ-RS, Loison et al., 2013) was used to evaluate participants’ motor imagery ability prior to take part to the psychophysical experiment. The average score (36.7±8.8; maximum score = 56) indicated good kinesthesis imagery ability. The protocol was approved by the clinical Ethics Committee of the University Hospital of Dijon (registered number 2009-A00646-51) and complied with the Declaration of Helsinki. Written informed consent was obtained from all participants. All of them were naïve as to the purpose of the experiment and were debriefed after the experimental session.

#### Experimental procedure

The aim of this experiment was to examine whether gravity constraints are integrated into mental movement simulation. One very simple and natural movement that is modulated by gravity and the biomechanics of the system is a free pendular movement. Previous studies showed that participants tend to adopt a pace that is faster (respectively slower) when gravity increases (respectively decreases) (Mechtcheriakov et al., 2002; White et al., 2008) or when the inertia of the system increases (Hatsopoulos and Warren, 1996) even if participants are instructed to maintain a constant pace. Here, we asked participants to actually perform free rhythmic movements in two different biomechanical conditions that predict different periods of motion. We then asked the participants to mentally simulate the same movements in these two conditions. We controlled that, as shown before, actual and spontaneous periods match and are modulated by simple biomechanical parameters. We reasoned that if biomechanical parameters, including gravity, are also relevant to motor imagery, the time taken to mentally simulate the same number of movement cycles in the two conditions should be different. We developed a computational model of the task (see supplementary materials) to support our prediction. The natural period depends on the inertia (I_i_) of the four different subsystems that compose a physical pendulum (upper arm, lower arm, wrist and loading condition of a hand-held bottle), total mass (m), gravity (g) and position of the center of mass (l) of the equivalent system according to:

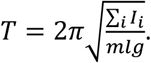

Participants were comfortably seated in an armless chair and maintained the dominant arm outstretched and vertically aligned with gravity. In the *light* loading condition, participants held an empty plastic bottle (0.039kg). In the *heavy* loading condition, participants held the same bottle, but full of water (1.089kg); therefore, increasing the inertia of the arm-bottle system. We positioned markers on the bottle, as well as on the shoulder, elbow, forearm, and wrist joints and recorded their 3D position (200Hz, dual-pass autoregressive filter at 20Hz) with a Vicon motion tracking system. Participants were asked to perform rhythmic movements at a free comfortable and natural pace for 30 seconds with the outstretched arm, holding either the empty bottle or the full bottle. Each bottle was used five times and trials were randomly interleaved. We then extracted the period of oscillations with a Fast Fourier Transform.

We repeated the same experiment in motor imagery. Participants adopted the same starting posture, but were now asked to mentally simulate 15 continuous cycles of movement in each loading condition (four with the empty bottle and four with the full bottle, randomized). The participants were instructed to keep their arm fixed along the body. As previously, they did not have an imposed pace to follow. However, they had to imagine performing constant movement oscillations amplitudes throughout the experimental session. The same experimenter triggered a stopwatch when the participant said ‘GO’ and stopped it at his/her ‘STOP’ injunction.

Finally, we pushed the exercise one step further and asked the participants to mentally simulate the same pendular movements, but in two unnatural environments. In the first, they had to imagine they were weightless (0g condition) and in the second, they had to project themselves in hyper-gravity (2g condition). Participants watched short movie clips of unrelated movements (jumps) performed in weightlessness and hyper-gravity contexts (recorded during parabolic flights). As before, there were four trials of 15 cycles in each gravitational condition randomly presented. The posture adopted was the same as in the imagined light condition.

#### Data analysis

Normal distribution of the variables was verified before using parametric statistical tests (Shapiro-Wilk W test; p>0.05). Periods were compared within each modality (*executed* and *imagined*) using paired t-tests. We also calculated linear correlation between the periods predicted by the model and the measured periods, separately in each modality, by taking into account the different anthropometric characteristics of participants. Data processing and statistical analyses were done using Matlab (The Mathworks, Chicago, IL).

## Results

Here, our main objective is to refine the roles of the anterior and posterior insulae with respect to the processing of gravity in the control of actions. We compared neuronal activations between executed and imagined hand movements in an MRI scanner. While both modalities require a cognitive step for movement preparation, only executed movements process feedback signals. We also developed a simple biomechanical model to demonstrate that even during motor imagery, biomechanical factors are integrated into mental states and thus influence the spontaneous timing of the mental action.

### Actual and mental movements activate different parts of the insula

Participants performed actual and mental movements of hand flexion-extension in the sagittal plane at free pace. Table 1 summarizes the brain network involved in the execution (*executed*>*rest*) and the mental simulation (*imagined*>*rest*) of hand movements. In actual movement production, the largest clusters were identified in the left primary motor and somatosensory cortices, in the bilateral supplementary motor area (SMA), as well as in the right cerebellum (lobes IV, V, VI). Activations were also highlighted in the left thalamus, in the left premotor cortex, in the left cerebellum (VI), and in the left middle temporal gyrus. For the imagined hand movements, the largest clusters were recorded in the right cerebellum (lobes VI) and the SMA bilaterally. The right inferior frontal gyrus, the left putamen, the right inferior frontal operculum, and the left inferior parietal lobule were also activated. Figure 1 depicts the main brain areas activated during actual movement production and its mental simulation.

**Table 1.** Significant activations to the contrasts [executed>rest] and [imagined>rest] (p-corrected for multiple comparisons<0.05). Brain lobe, regions from AAL atlas and coordinates (x, y, z) in the MNI-space are reported. The first region has the highest number of voxels in the cluster and the other regions [between brackets] belong to the cluster with lower number of voxels. The two last columns correspond respectively to the maximum T-value and the number of voxels in the cluster.

| **Lobes** | **X** | **Y** | **Z** | **T-value** | **Voxels**<br>**in** |
| --- | --- | --- | --- | --- | --- |
| *AAL Regions* |  |  |  |  |  |
| *Executed>Rest* |  |  |  |  |  |
| *Parietal, frontal, sub-lobar and limbic lobe, Left postcentral gyrus, [Left precentral gyrus, Left superior motor area, Left midcingulate area, Left supramarginal, Left rolandic operculum, Left inferior parietal lobule, Right superior motor area]* | -30 | -26 | 56 | 17.57 | 3260 |
| *Anterior lobe of cerebellum, Right cerebellum (lobules IV, V, VI), [Vermis IV, V, VI]* | 22 | -50 | -22 | 15.80 | 1738 |
| *Parietal lobe, Left thalamus* | -14 | -22 | 6 | 9.99 | 75 |
| *Sub lobar & frontal lobe, Left insula, [Left rolandic operculum]* | -44 | 0 | 2 | 9.32 | 208 |
| *Parietal lobe* | 56 | -32 | 24 | 9.22 | 233 |
| *Right supramarginal* |  |  |  |  |  |
| *Frontal lobe, Left precentral gyrus* | -56 | 4 | 32 | 7.96 | 30 |
| *Anterior lobe of cerebellum, Left cerebellum (lobules VI)* | -30 | -54 | -26 | 7.88 | 57 |
| *Sub lobar, frontal and temporal lobe, Right rolandic operculum* | 46 | 2 | 6 | 7.82 | 103 |
| *[Right inferior frontal operculum], [Right insula]* |  |  |  |  |  |
| *Temporal lobe, Left middle temporal gyrus* | -52 | -70 | 6 | 7.78 | 20 |
| *Anterior lobe of cerebellum, Vermis IV, V* | 2 | -44 | -2 | 7.59 | 17 |
| *AAL Regions* |  |  |  |  |  |
| *Imagined>Rest* |  |  |  |  |  |
| *Anterior lobe of cerebellum, Right cerebellum (lobules VI)* | *34* | *-54* | *-28* | *10.79* | *138* |
| *Sub lobar, frontal and temporal lobe, Left rolandic operculum* | *-54* | *6* | *10* | *10.15* | *356* |
| *[Left inferior frontal operculum], [Left insula]* |  |  |  |  |  |
| *Frontal lobe, Left superior motor area, [Right superior motor area]* | *-6* | *-8* | *66* | *9.04* | *125* |
| *Parietal lobe, Left supramarginal* | *-48* | *-36* | *22* | *8.47* | *88* |
| *Frontal lobe, Right inferior frontal gyrus* | *46* | *42* | *-6* | *8.28* | *49* |
| *Sub-lobar, Left putamen* | *-26* | *-4* | *12* | *8.24* | *27* |
| *Frontal lobe, Right inferior frontal operculum* | *56* | *8* | *20* | *7.77* | *21* |
| *Parietal lobe, Right supramarginal* | *60* | *-32* | *26* | *7.69* | *44* |
| *Parietal lobe, Left inferior parietal lobule* | *-50* | *46* | *40* | *7.59* | *12* |
| *Sub lobar, frontal and temporal lobe, Right rolandic operculum* | *50* | *6* | *6* | *7.23* | *27* |
| *Frontal lobe, Right inferior frontal operculum* | *54* | *16* | *14* | *6.80* | *10* |

As the main objective of this report was to investigate whether the insula, known to be involved in the processing of gravity-relevant signals during executed movements, was also activated during imagined hand movements, we further focussed our analysis on that brain region. The contrasts *executed*>*rest* and *imagined*>*rest* revealed the activation of both the right and left insulae. The analysis did not highlight clear differences in insular responses between the two contrasts, likely due to the size of the clusters covering the insula. Noteworthy, the peak activity was localized in a more anterior part of the insular cortex in imagined movements (compare Figure 1A and 1B). To further compare the activation of the insular cortex between executed and imagined movements, we examined the contrasts executed-rest>imagined-rest and imagined-rest>executed-rest by using an anatomical mask of the bilateral insula (p<0.05, corrected for multiple comparisons). Interestingly, we identified one cluster in the posterior part of the left insular cortex (x=−38, y=−26, z=24) when contrasting executed with imagined movements (Figure 2), but not when we contrasted imagined with executed hand movements.

**Figure 2.**
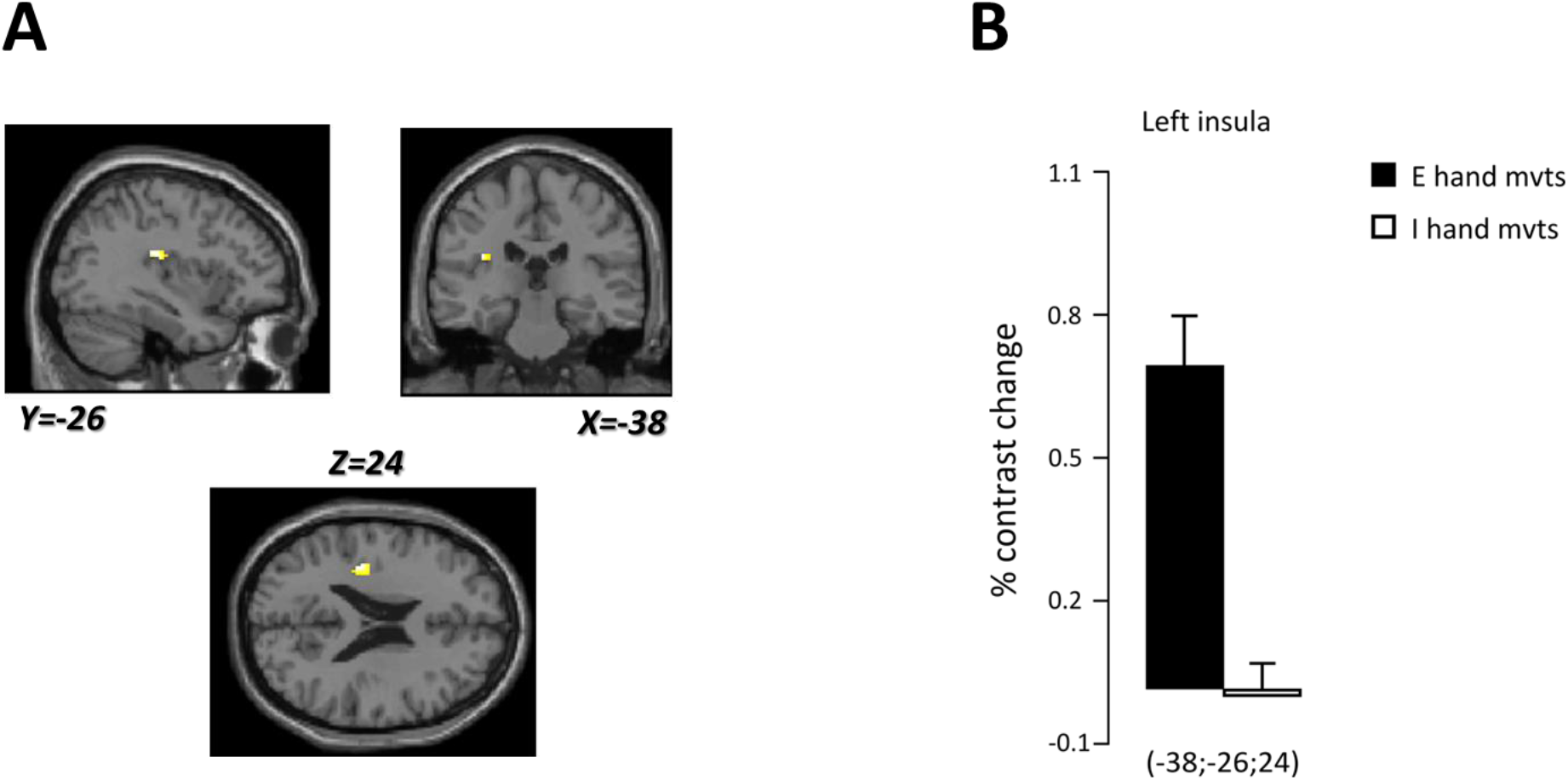
A. Localization of the cluster found in the left insular cortex (yellow) specifically engaged during executed compared to imagined hand movements. Brain responses are projected onto the sagittal (Y=−26), coronal (X=−38) and axial (Z=24) views of the T1-single subject template of MNI. B. Percent contrast change in the left insular cortex during executed (black) and imagined (white) vertical hand movements.

### Imagined movements integrate gravito-inertial constraints

The aim of the psychophysical experiment was to test whether gravito-inertial constraints are included into the mental movement simulation process. Participants imagined and performed rhythmic pendular movements in two inertial conditions in the lab environment and imagined the same movements as if they were in microgravity (0g) and hyper-gravity (2g).

Figure 3 (left panel) depicts the average periods spontaneously adopted by participants (in the normal gravity environment of the lab) in the *light* (empty bottle) and *heavy* (filled bottle) loading conditions and separately for the real (red bars) and imagined movement conditions (blue bars). The ANOVA revealed a main effect of *loading* condition (F_1,14_=14.62, p=0.002, η^2^=0.51), but no effect of *movement* condition (F_1,14_=0.02, p=0.90) or interaction effect (F_1,14_=0.01, p=0.74). Consistently, participants moved at a slower pace (i.e., spontaneously adopted longer periods) in the executed and imagined *heavy* movement conditions compared to the *light* loading condition. A correlation analysis within each loading condition and between both movement modalities confirmed these results. The correlation coefficient was 0.66 (p<0.001) between the executed and imagined light condition and 0.75 (p<0001) between the executed and imagined heavy condition. The model also predicts small changes with body segment lengths. To quantify this, we calculated the correlation between model predictions of the spontaneous period (by specifying the exact lengths of the upper arm, lower arm and wrist) and the measured period. We found significant correlations in the executed modality (r=0.54, p<0.001) and imagined modality (r=0.42, p=0.038).

**Figure 3.**
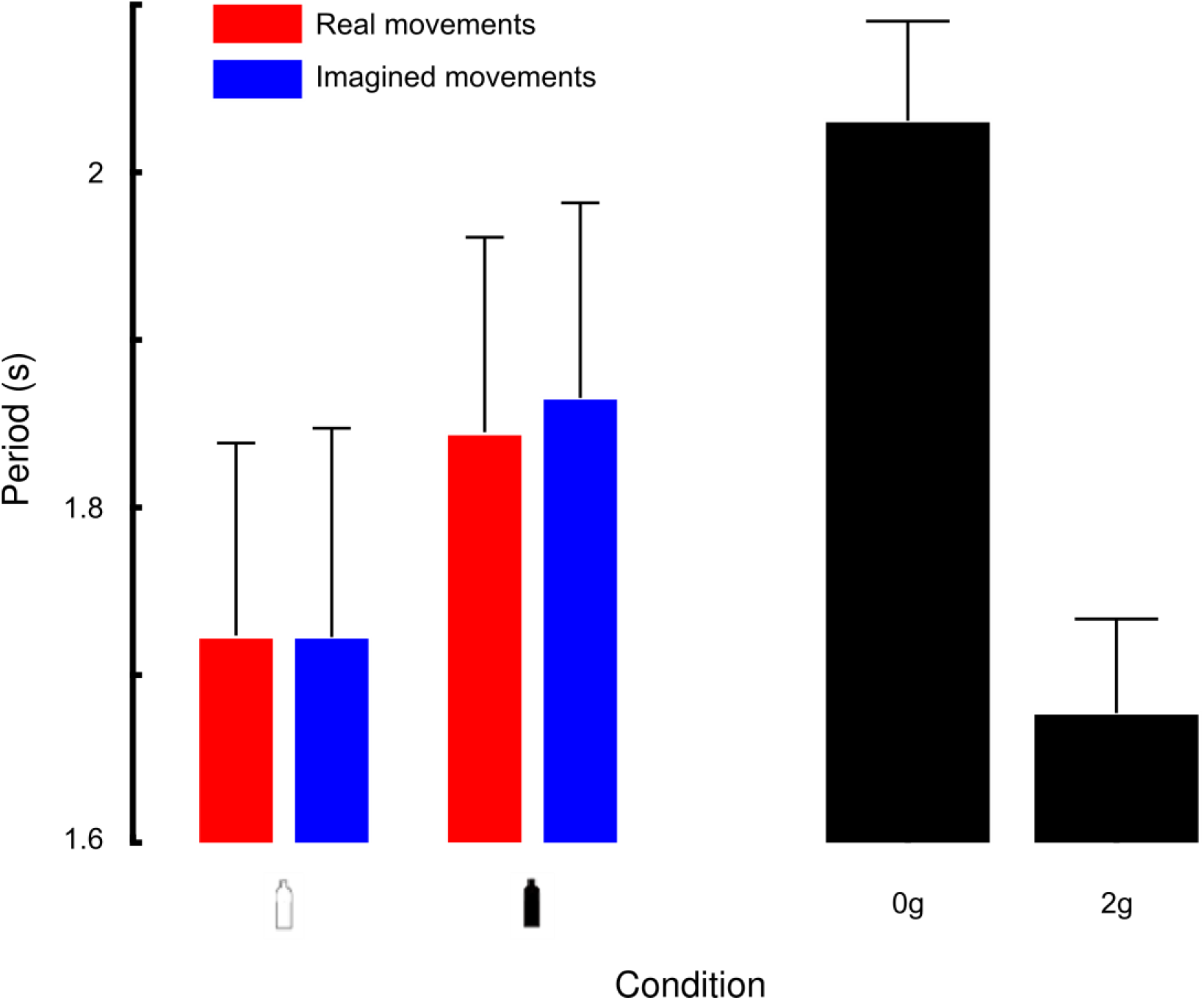
Mean periods of movements. (A) Spontaneous periods of movements in the light loading condition (open bottle) and in the heavy loading condition (closed bottle). Red and blue bars denote real and imagined movement conditions, respectively. (B) Periods measured during imagined movements performed in weightlessness (0g) or in hypergravity (2g). Error bars are SD.

We also indirectly tested the effect of the gravitational environment in which we mentally simulate the action. Remarkably, Figure 3 (right panel) shows that periods naturally adopted to mentally simulate the same task in weightlessness increased by 30% compared to hyper-gravity (t_14_=2.5, p=0.023, η^2^=0.23). Furthermore, one can also notice that periods in the 1g condition were longer than periods in the 2g condition (compare Fig. 3A-B). These effects are compatible with our model which predicts that movement become slower (and have a larger period) as gravity level decreases and conversely in hyper-gravity condition. Therefore, by assuming one can reliably imagine a movement in the normal Earth environment (1g), we used the model to predict what should be the theoretical gravitational level to account for this period change. To do so, we calculated, for each participant, the difference between periods (Δ*T*) in both conditions. We also personalized the biomechanical factors (m, l and J) to each participant in the model prediction. We then solved the following equation for alpha, which we interpret as a gravitational gain:

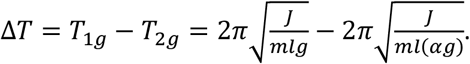

After algebraic development, one can easily find:

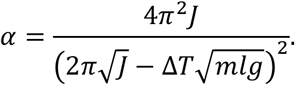

We found an average value of *α* = 1.79 (SEM=0.37), which is not exactly 2 but approaches a reasonable hyper-gravity level. A similar development allows to infer the estimated (imagined) value of weightlessness, using the above equation with an addition in its denominator:

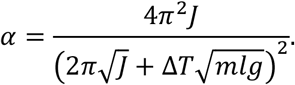

The average values of *α* are 0.61 (SEM=0.12), which is below 1 but clearly not zero. This is not surprising since the model is ill-defined for g=0 (it predicts infinite periods). It was shown previously that other neural mechanisms take over when g approaches zero (White et al., 2008).

Together, this experiment shows that participants do not only integrate biomechanical factors to simulate a movement including their own morphology, but also environmental factors such as gravity. This underlines a good capacity of kinaesthetic mental imagery and puts us on safe grounds to assume that gravity is indeed encoded in mental simulation.

## Discussion

In the present study our aim was to isolate the role of the anterior and posterior insulae in the processing of gravity by contrasting actual and mental movements. We found that mental movements activated brain areas similar to actual movements. This confirms that motor execution and motor imagery engage common neural representations (for a review, see Hardwick et al., 2018; Hétu et al., 2013). Our findings also revealed brain responses in the insular cortex in both actual and imagined movements. More specifically, we found posterior activation of the insula in movement execution and anterior activation of the insula in mental movement simulation. The gravitational force significantly alters motion dynamics during the execution of movements. The insula, known to process the effects of gravity via a stored internal representation of gravitational acceleration, is activated for actual hand movements (Indovina et al., 2005; Lacquaniti et al., 2014; Rousseau et al., 2016). Here, we made a step forward by showing that the insula is also activated when the task mainly relied on mental predictive states.

Visual tasks have been shown to activate the insula. In an elegant study, Lacquaniti and colleagues, demonstrated that an internal representation of gravity is activated by visual motion that appears to be coherent with natural gravity (Indovina et al., 2005; Lacquaniti et al., 2014). The insular activation associated to neural predictive mechanisms was also observed during experiment of visual motion when comparing prediction versus perception of visual motion (Cheong et al., 2012). Here, we showed that the insular cortex is activated during mental movement simulation in absence of gravity-relevant visual feedback. In line with our results, global body movements, such as active balance simulation task (Karim et al., 2014) and mental imagery of balance (Jahn et al., 2004; Malouin et al., 2003) elicited fMRI responses in the insula. Further, Sacco and colleagues showed that the insula plays an important role in mental imagery of tactile and proprioceptive sensations and therefore in the imagery of actions and sensations in general (Dijkerman et al., 2007; Sacco et al., 2006). Our results go one step further and demonstrate that the insular cortex integrates dynamical constraints – including gravity – to implement predictive mechanisms, even in absence of physical interaction between the body and the environment.

The posterior insula was more activated during actual hand movements compared to imagined hand movements, which, to the best of our knowledge, is an original finding. The largest number of afferences with the sensorimotor system in the posterior insula may explain why its activity during real movements was higher compared to the anterior insula. At this stage of investigation, we can speculate that the anterior and posterior insulae possess different roles. On the one hand, the anterior insula may implement predictive mechanisms (internal simulations) to integrate the expected dynamical constraints caused by gravity when no gravity-relevant feedback is available. On the other hand, the posterior insula could be fundamental to process the effects of gravity by including the information conveyed by sensory feedback.

We also examined the temporal features of actual and mental movements performed under different gravito-inertial constraints. Our results showed, regardless of task dynamics, a high temporal correspondence between them. This isochronism extends previous findings that reported comparable durations of actual and mental movements (Gentili et al., 2004; Guillot and Collet, 2005; Papaxanthis et al., 2012) and reinforces the well-documented idea of a similar neurocognitive network between actual and mental states (Grosprêtre et al., 2019; Ruffino et al., 2017). Temporal similarities between actual and mental movements reinforce the idea that the brain preserves an accurate internal model of arm and environmental dynamics. A controller and a forward internal model, integrating task dynamics, could provide similar durations for actual and mental movements. The controller would generate the appropriate neural commands necessary to move the segment into the vertical plane. The forward model would relate the sensory signals of the actual state of the segment (e.g. position, velocity) to the motor commands, and would predict its future states. In the case of imagined movements, the motor commands do not reach muscles, i.e., no movement occurs. However, a copy of these motor commands is still available to the forward model, which predicts the future states of the segment and therefore provides temporal information very similar to that of actual movements. In general, our results indicate that the CNS maintains an accurate representation of gravito-inertial dynamics and uses this representation to appropriately simulate arm movements in the vertical plane.

Strikingly, albeit participants were in a normal gravity environment, they could mentally simulate arm movements in radically new contexts (i.e., microgravity or hyper-gravity) by appropriately changing their temporal features (see modelling results). Specifically, movement imagined in microgravity and hyper-gravity were, correspondingly, slower and faster than movements imagined in normal gravity. This further suggests that the brain can imagine movements in a purely cognitive process (none of the participants had a personal sensorimotor experience of these altered contexts) in other environments than the habitual Earth’s environment. Should the brain had imagined movements irrespective of gravitational constraints, similar durations would have been observed between the different conditions. Note that the integration of gravity into motor prediction per se is not a *de facto* condition. Indeed, it has been proposed that mental imagery of objects’ visual motion does not have access to the internal model of Earth gravity, but resorts to a simulation of visual motion compatible with a 0g environment (Gravano et al., 2017). It is possible that when trying to extrapolate object trajectories in the environment, the internal model of gravity for these predictive tasks is triggered by visual information of the object motion. In contrast, during internally generated predictive movements (mental imagery), the brain could rely on a more physical (in the Newtonian sense) internal model of gravity (Mcintyre et al., 2001). The question of whether the body is actively involved in the task may therefore be decisive as to which gravitational representation to load. It remains to be tested, however, which part of the insula and therefore which modality of the gravitational model is activated when purely imagining an object moving in the gravitational field as opposed to motor imagery.

## Conclusion

Before executing a movement, the CNS generates a motor command (efference copy). Neural mechanisms using the currently issued efference copy simulate the upcoming movement and its sensory consequences (Wolpert and Kawato, 1998). This is the role of the forward model (Figure 4). This active mental process integrates, among other parameters, the dynamical effects induced by the gravitational field. In case of errors, feedback signals available after various delays will be used to update both the forward model and the controller. Neural correlates of this forward model include the cerebellum (Diedrichsen et al., 2005) and the contralateral and supplementary motor area (Bursztyn et al., 2006).

**Figure 4.**
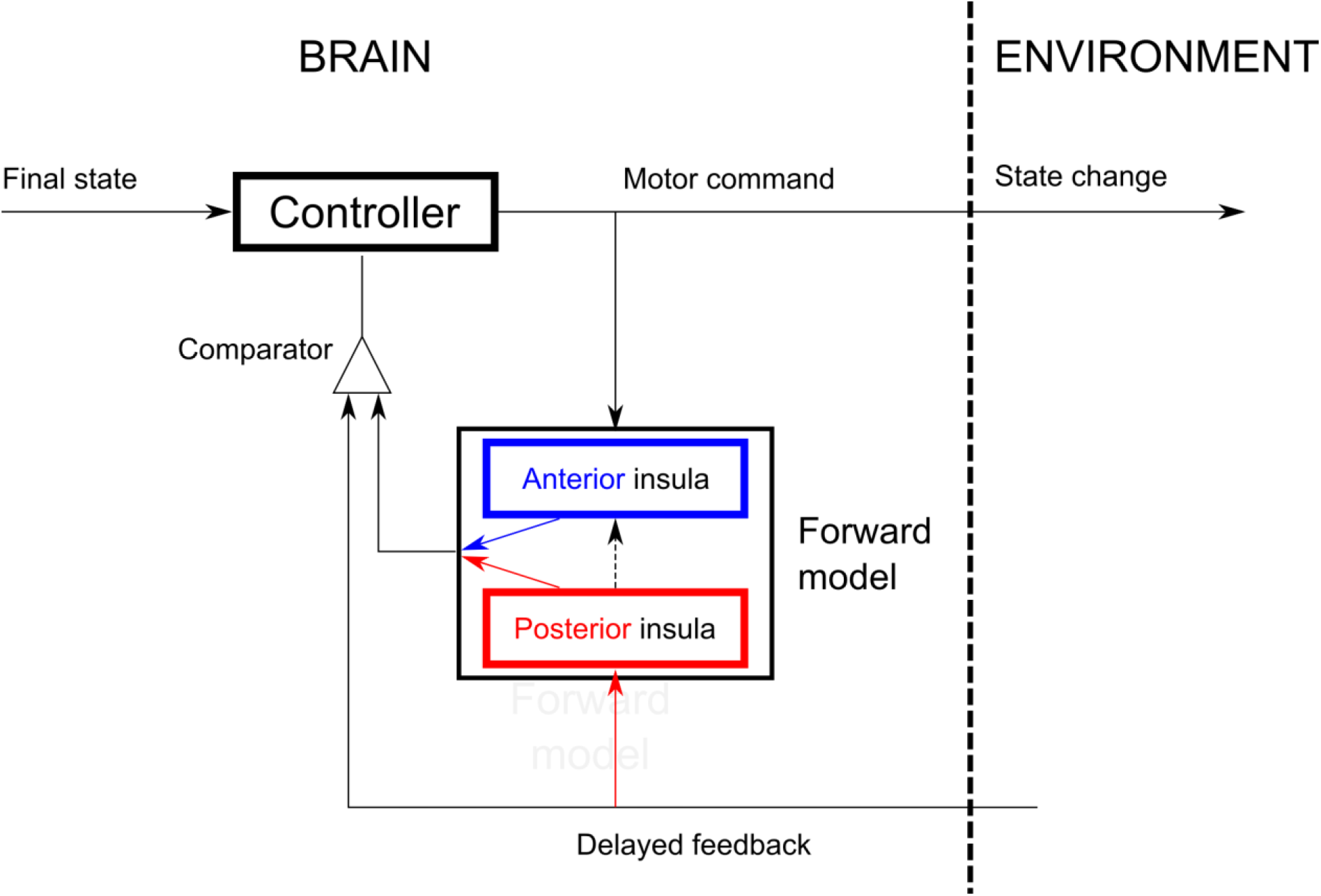
Conceptual sketch illustrating the different roles of the posterior and anterior insula in internal models, with respect to the specific processing of gravity.

Our data suggest there are two complementary representations of gravity in the insular cortex, conceptually illustrated in Figure 4. The first, situated in the posterior insula (red box), is used when sensory feedback is available. In that case, feedback signals directly update the internal representation of gravity (vertical red arrow). The output of the forward model (slanted red arrow) is compared with the feedback to evaluate the presence of errors. The second, offline, representation of gravity is held in the anterior part of the insula (blue box) and can be considered as a proxy of the first. Since no task-relevant feedback is available, we posit this dual internal model of gravity cannot be adapted during motor imagery, but is accessible for simulations. In this case, the posterior insula is not included in the online control loop. During motor imagery, the output of the anterior insula (slanted blue arrow) is compared with feedback. These two regions are unidirectionally connected in such a way to allow the posterior insula to update the abstract representation of gravity in the anterior insula (vertical black dashed line). The question remains open as to why two different versions exists?

## Acknowledgments

The authors declare no conflict of interest. This research was supported by the « Conseil Général de Bourgogne-Franche Comté » (France), the « Fonds Européen de Développement Régional » (FEDER) and the « Centre National d’Etudes Spatiales » (CNES), and the ANR (Motion, ANR-14-CE30-0007). The authors also thank J. Gaveau for interesting discussions to refine the psychophysics experiment.

## Supplementary material

### Modelling the upper limb and an object as a physical compound pendulum

The aim of this technical appendix is to find the preferred period of a physical pendulum that models the system composed by the upper arm, the forearm, the hand and a light bottle containing water. The segments are considered collinear and joints are rigid (Fig. 1).

**Figure S1.**
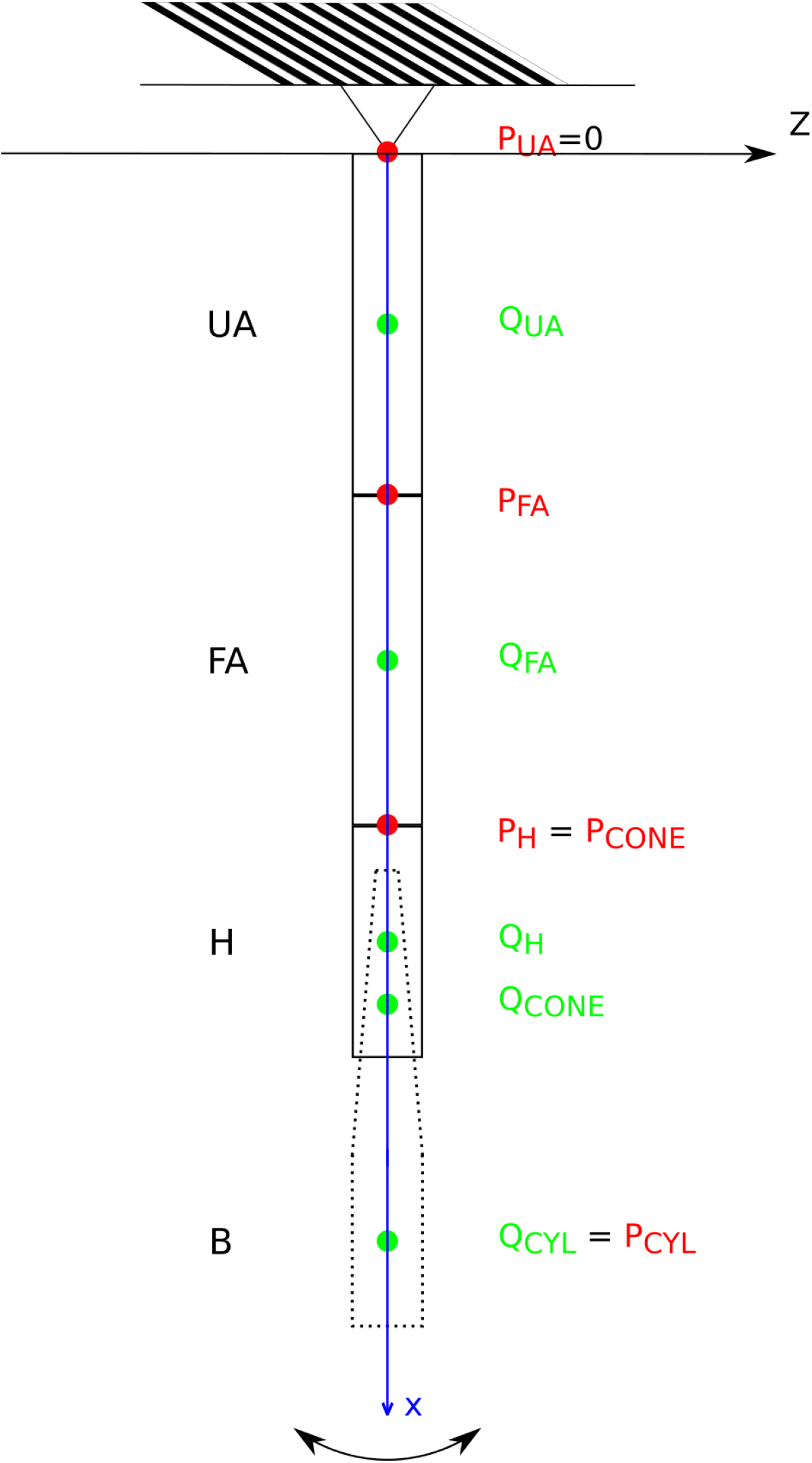
System model and equivalent mathematical pendulum.

### 1. Background

Let’s consider the plane motion of a material body suspended on a pivot point (point O in supplementary Fig. 1) and allowed to rotate freely about z. A right-handed Cartesian coordinate system is centred on the pivot point O and ϕ denotes the amplitude of oscillations. The equation of motion in moment terms for such mechanical system is given by:

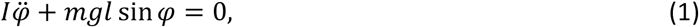

where *I* is the moment of inertia of the system about z, and *m* and *l* are the total mass and position of its center of mass (g is gravitational attraction, *g* = 9.81 ms^−2^). The equation above can be reduced to

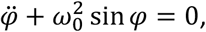

with

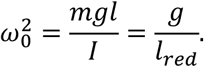

The last expression corresponds to the equation of a mathematical pendulum of equivalent length *l*_*red*_ and submitted to gravity. Notice that it is independent of mass. The natural (resonant) period, under the assumption of small oscillations can be obtained by:

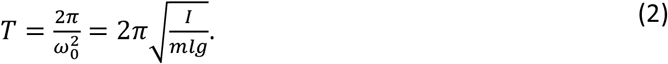

For completeness, note that adding a damping coefficient proportional to angular velocity in Eq. 1 does not influence natural period. We need to calculate the period *T* for a 4-part physical compound pendulum:

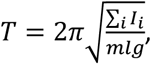

where I is replaced with the sum of moment of inertia of each system part, calculated with respect to the same reference (y axis).

### 2. Calculation of individual *I_i_*

As illustrated in the previous section, the compound physical pendulum can be decomposed in several parts (Fig. 1). In this section, we calculate the moment of inertia, the mass, and position of centre of mass of each subsystem.

#### 2.1 Inertia of biological parts (upper arm, forearm and hand)

While these three components can be modelled by three different uniform rods of different masses and physical dimensions, we use more accurate tabulated values instead (mass is not uniformly distributed). The moment of inertia *I* of a mass *m* about an axis of rotation is *I*_0_ = *mr*^2^ (*r* is the distance from the point mass to the axis of rotation). Using the parallel axis theorem, we can express moment of inertia with respect to another (parallel) axis of rotation situated at a distance *d* from the first:

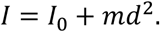

The equation above can be transformed to explicitly use the radius of gyration *k*:

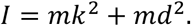

Anthropometric data about biological segments are available in tables (Winter, 2009). Mass and length are expressed in proportion (%) to total body mass and total body height, respectively. Similarly, the position of the center of mass (*C*) and radius of gyration (*K*) are expressed in proportion to segment length. For the upper arm, the moment of inertia with respect to the proximal part of the segment (close to the shoulder) is

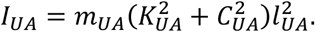

We get the same expression for the forearm and the hand:

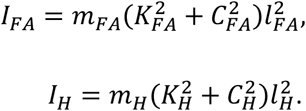

In the three equations above, *m* and *l* are the mass and length of each segment and *K* is the radius of the segment with respect to its center of mass. These values are tabulated and can be individualized from individual subject body height (H) and mass (M). The table below summarizes the values of each parameter for the lower arm, the forearm and the hand:

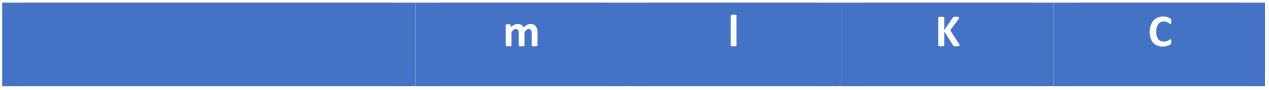
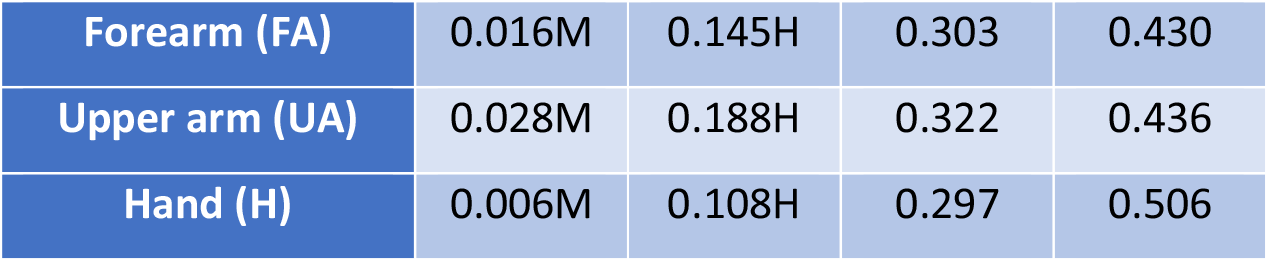

#### 2.2 Inertia of the passive system

The experimental context comprised two conditions. In condition 1, the upper arm-forearm-hand system was outstretched and the hand held the empty bottle that was oriented along the segment main axis. Condition 2 was identical to condition 1 except that the bottle contained water.

##### Conditions 1

Since the mass of the empty bottle is very light (0.039kg) compared to the mass of the biological part of the system, we neglect the additional inertia induced by the bottle.

##### Condition 2

The bottle (height *h*_*b*_) can be modelled by a cylinder (CYL) of radius *R* topped with a frustum cone (CONE) with small radius *r* as shown in Figure 2 below.

**Figure S2.**
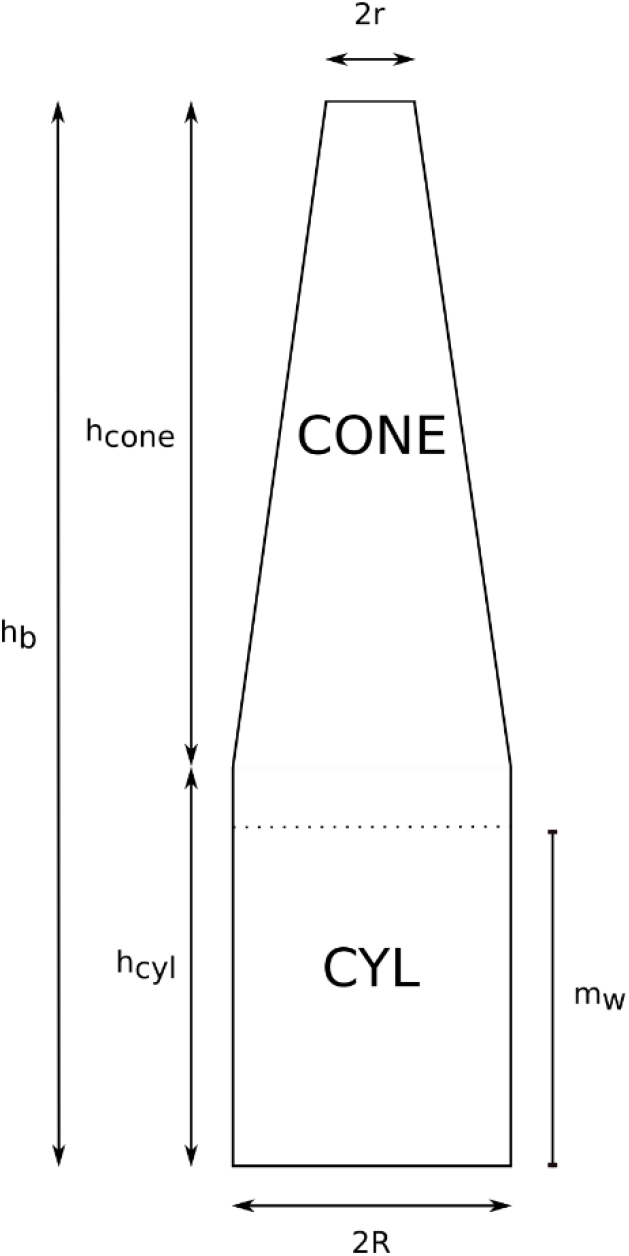
Model of the bottle.

For the sake of the calculation of moments of inertia, we will consider each part of the bottle separately (CYL and CONE). The mass concentrated in the CYL part of the bottle and on a height of water *n*_*w*_ is *m*_*CYL*_ = 0.564 kg. By symmetry, the position of the center of mass of such solid is halfway its height 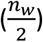. We now need to calculate the moment of inertia of the CYL about an axis through the center of mass but different from its main axis. The moment of inertia of a cylinder about an axis passing through its diameter and its center of mass is:

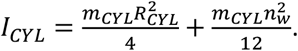

The conic part of the bottle is also full of water (*m*_*CONE*_ = 0.525 er ()). The problem to be solved consists in finding the position of the center of mass of a frustum cone and its moment of inertia
about an axis perpendicular to its longitudinal axis. Figure 3 illustrates the problem in 2 dimensions and reports all relevant parameters. We will use variational calculus to compute the position of the center of mass of this solid and its moment of inertia.

**Figure S3.**
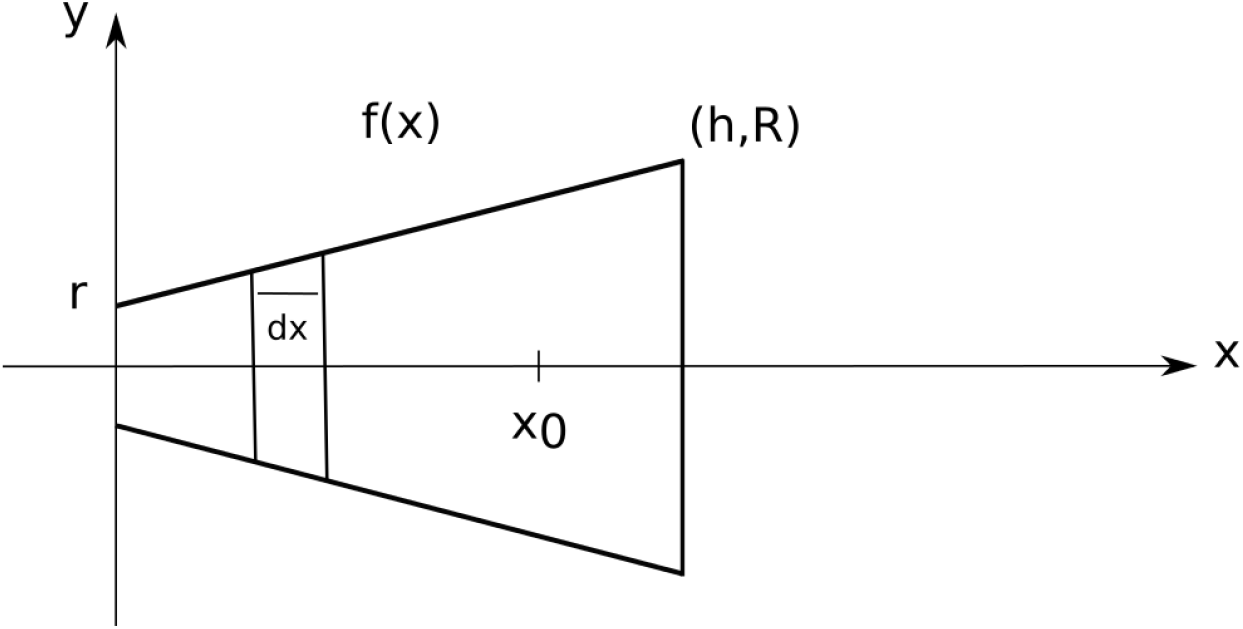
Frustum cone.

We first find the equation of the slant of this shape as a function *f* of *x*:

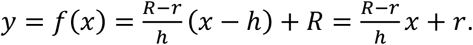

The position of the center of mass *x*_0_ of this solid is given by:

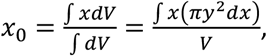

where the infinitesimal volume element *dV* is the volume of a thin disk of thickness *dx* and radius *y*. We know that the volume of a frustum cone is:

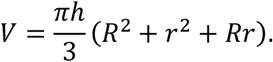

The resolution of the integral in the numerator of Eq. X yields:

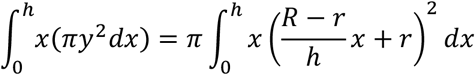

After simplification with the total volume, we find:

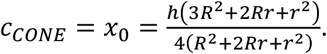

Notice that this expression simplifies to 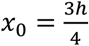 for a cone (*r* = 0).

Having the center of mass of a frustum cone, one can now calculate its moment of inertia following the same logic. Let us first calculate the moment of inertia *I*_*x*_ of the solid about the x-axis (see Figure 3). Since we know that the moment of inertia of a thin disk (mass *dm*, radius *y*) with respect to the x-axis is 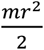, we have:

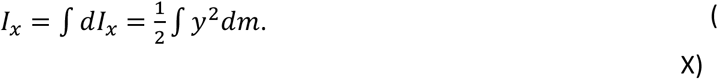

Since the volume and mass are related by density through *dm* = *ρdV* = *ρπy*^2^*dx*, we have:

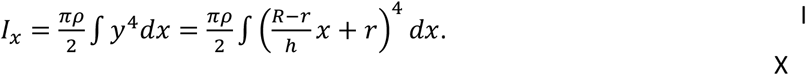

Let us now calculate 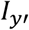, where *y’* is an axis parallel to the y axis passing through a diameter of the disk. For that solid, we know that (and using Eq. X):

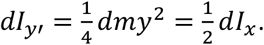

By application of the parallel axis theorem, we can now deduce:

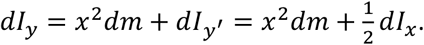

Therefore:

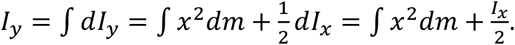

By specifying the boundaries of the domain of integration, we have to calculate:

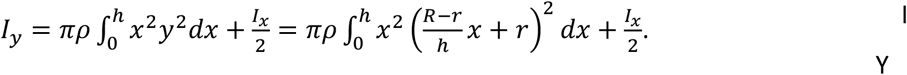

Both terms of the above equation are easily calculated. After development and simplification, we find first the first term of the expression above:

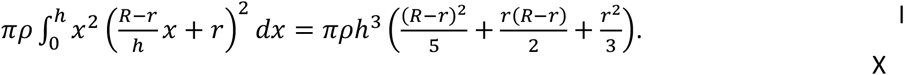

We then derive the value of *I*_*x*_:

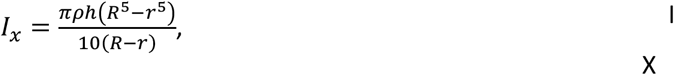

and plug it back into *I_y_* which yields the value we are looking for:

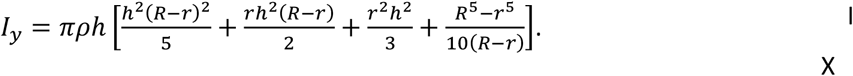

### 3. Considering the whole compound system

To sum up, we have calculated the individual positions of center of mass, masses and moments of inertia of each component relative to their center of mass or to a specific position:

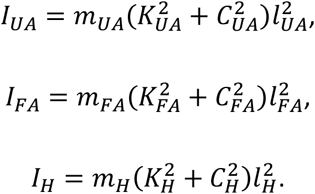

The above values are common to both conditions. In condition 2, the dynamics of the off-centered mass is also considered:

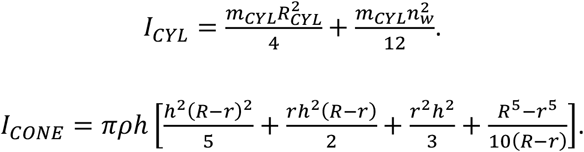

The total moment of inertia of the compound system is the sum of the individual moments. However, all the *I*_*i*_ were not calculated with respect to the same reference. Therefore, before summing up *I*_*i*_, we must apply the parallel axis theorem to derive the moment of inertia about the pivot point (elbow joint, about the y axis, Fig. 1). We label these converted moments of inertia *J*_*i*_. The sketch in Figure 1 summarizes the problem and reports the position about which *I*_*i*_ were calculated (*P*_*i*_) and the centers of mass (*q*_*i*_) with respect to the origin of the reference frame.

We get:

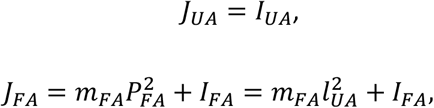

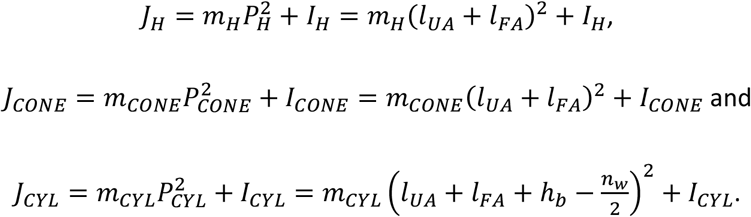

Finally,

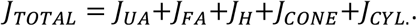

The objective of this development was to find the natural period of the system:

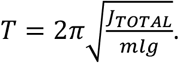

The total mass of the system is simply the sum of the mass of its components, 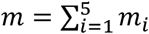. The length of the system center of mass is the weighted sum of the individual masses (*q*_*i*_ are distances to each center of mass, from the pivot point):

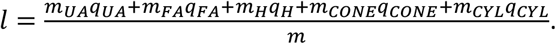

The natural period of the system varies between conditions. Larger inertias contribute to increase the period while a larger mass and a more off-centred position contribute to decrease period. We have fixated the anthropometric values to typical values (H=1.7m and M=70kg). However, in the analyses, we took into account (subtle) individual subject characteristics in terms of segment lengths. The figure below plots the simulated natural periods for the two conditions and clearly reveals that the period should increase if the bottle is full of water.

**Figure S4.**
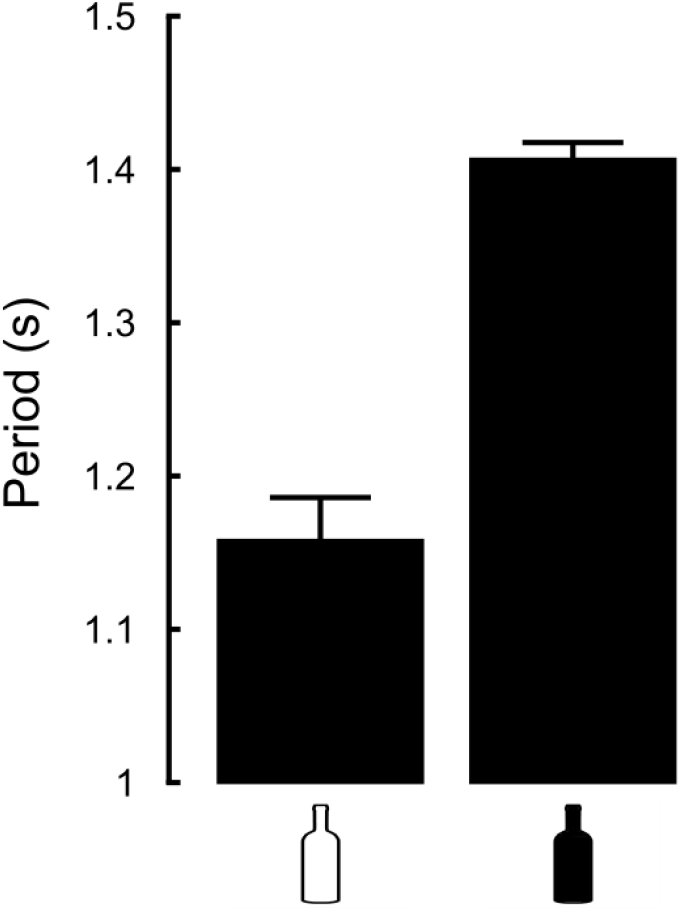
Simulated periods in both conditions (open symbol = empty bottle; closed symbol = full bottle). SD correspond to standard deviation across subjects (n=15) that reflect small differences in their individual biomechanics.

